# The neuron-intrinsic membrane skeleton is required for motor neuron integrity throughout lifespan

**DOI:** 10.1101/2025.02.23.639536

**Authors:** Carrie Ann Davison, Daniela Garcia, Chengye Feng, Hongyan Hao, Erik M. Jorgensen, Marc Hammarlund

## Abstract

Axons experience physical stress throughout an organism’s lifetime, and disruptions in axonal integrity are hallmarks of both neurodegenerative diseases and traumatic injuries. The spectrin-based membrane periodic skeleton (MPS) is proposed to have a crucial role in maintaining axonal strength, flexibility, and resilience. To investigate the importance of the intrinsic MPS for GABAergic motor neuron integrity in *C. elegans*, we employed the auxin-inducible degron system to degrade β-spectrin/UNC-70 in a cell-type specific and time-dependent manner. Degradation of β-spectrin from all neurons beginning at larval development resulted in widespread axon breakage and regeneration in VD/DD GABAergic motor neurons in both larval and adult animals. Similarly, targeted degradation of β-spectrin in GABA neurons alone resulted in extensive breakage. Moreover, we found that depleting β-spectrin from the mature nervous system also induced axon breaks. By contrast, epidermal β-spectrin was not required for axon integrity of VD/DD neurons. These findings demonstrate the cell-intrinsic importance of neuronal β-spectrin/UNC-70 for axon integrity both during development and in adulthood.

## Introduction

Throughout the lifetime of an organism, neurons are subject to physical stress. Traumatic injury can disrupt axon integrity; even without trauma, axons must maintain resilience for the life of the neuron. The spectrin-based membrane periodic skeleton (MPS) has emerged as a key support for axon integrity. The MPS is a lattice framework of actin rings connected by spectrin tetramers (D’Este et al., 2015; He et al., 2016; Xu et al., 2013) that is proposed to provide mechanical resistance and flexibility to neurites under physiological strain (Hammarlund et al., 2007; Krieg et al., 2014). In axons, the MPS exhibits a periodicity of approximately 190 nm (D’Este et al., 2015; He et al., 2016; Xu et al., 2013), and this periodicity is important for proper neuronal function including regulating axon diameter and conduction (Costa et al., 2020), organizing discrete membrane domains (C. Zhang et al., 2013), regulating endocytosis in the axon-initial segment (Wernert et al., 2024), promoting axonal transport (Jia et al., 2019; Lorenzo et al., 2019), and limiting tension propagation to enable localized signaling (Català-Castro et al., 2025). Disruptions in the MPS cause axon degeneration in *C. elegans* (Hammarlund et al., 2007; Krieg et al., 2014, 2017), *Drosophila* (Hülsmeier et al., 2007; Lorenzo et al., 2010), and mice (Huang, Zhang, Ho, et al., 2017; Liu et al., 2020; Lorenzo et al., 2019). In humans, mutations in α and β spectrin cause a variety of neurological disorders including spinocerebellar ataxia type 5 (Ikeda et al., 2006), early infantile epileptic encephalopathy-5, as well as other forms of developmental delay and intellectual disability (reviewed in (Lorenzo et al., 2023)). A deeper understanding of the function of the MPS within neurons is crucial for advancing nervous system resilience in injury and disease.

Spectrin tetramers consist of two α and two β subunits each containing multiple spectrin repeat domains that fold into three-helix bundles (Speicher & Marchesi, 1984; Yan et al., 1993). These bundles can reversibly unfold to act as spring-like “shock absorbers” and buffer tension in axons (Dubey et al., 2020). While humans have two α-spectrins and five β-spectrins, invertebrates have only one of each. In *C. elegans*, the sole β-spectrin is encoded by *unc-70* (Hammarlund et al., 2000); for simplicity, we will typically refer to this protein simply as β-spectrin.

β-spectrin has been proposed to provide tension and mechanical resistance to axons (Hammarlund et al., 2007; Krieg et al., 2014, 2017). Loss of β-spectrin causes spontaneous breakage of sensory and motor neurons due to the mechanical stress of body movement (Hammarlund et al., 2007; Krieg et al., 2014). Some evidence suggests this is due to a cell-intrinsic function for the spectrin cytoskeleton. For example, GABA neurons express β-spectrin (Taylor et al., 2021), and the endogenously tagged MPS has been visualized in *C. elegans* neurons (Glomb et al., 2023; He et al., 2016; Krieg et al., 2017), consistent with MPS function in neurons. Further, cultured *C. elegans* touch neurons have reduced apparent membrane tension in the absence of β-spectrin, indicating that loss of β-spectrin alters the cell-intrinsic properties of the neuron (Krieg et al., 2014).

However, β-spectrin is also expressed in other tissues including the epidermal tissues surrounding neurons, (Coakley et al., 2022; C. Wang et al., 2020; White et al., 1997). The importance of the spectrin cytoskeleton in surrounding tissues for axon function and integrity has been increasingly appreciated. For example, in mammals, loss of αII and βII spectrin in Schwann cells disrupts myelination of peripheral nervous system (PNS) axons (Susuki et al., 2011), and loss of glial βII-spectrin disrupts paranode formation and maintenance (Susuki et al., 2018). In *C. elegans,* epidermal β-spectrin has been shown to contribute to the axon integrity of touch receptor neurons (TRNs) (Bonacossa-Pereira et al., 2022; Coakley et al., 2020, 2022; Das et al., 2021; Krieg et al., 2014, 2017). The TRNs are embedded within the epidermis (Chalfie & Sulston, 1981), and the attachment between the neuron and epidermis has been shown to protect axons from movement-induced damage (Bonacossa-Pereira et al., 2022; Coakley et al., 2020).

Whether the spectrin cytoskeleton acts in neurons or in associated tissue to preserve axon integrity in neuron types that are not embedded in skin is unclear. The *C. elegans* DD and VD GABA neurons extend commissural axons that contact the skin, but are not embedded in it (Sulston & Brenner, 1997; White et al., 1997). β-spectrin in both muscle and the epidermis has been shown to induce DD/VD axon defects in response to hypergravity (Kalichamy et al., 2020). Furthermore, it has been reported that loss of β-spectrin from the epidermis results in breakage of DD/VD axon commissures, while loss from neurons has no effect (Coakley et al., 2022). On the other hand, a neuron-intrinsic function remains a compelling hypothesis based on the structure of the MPS (Xu et al., 2013) and on data from neurons in culture (Das et al., 2021; Dubey et al., 2020; Krieg et al., 2014). Some previous studies investigating the importance of β-spectrin for the integrity of the DD and VD GABA neurons have used full body mutations in *unc-70,* and did not fully address the question of the relative contribution of intrinsic and extrinsic spectrin to axon integrity (Hammarlund et al., 2007; Kalichamy et al., 2020). Thus, the importance of intrinsic and extrinsic spectrin for axon integrity in DD and VD GABA neurons remains unclear.

To characterize the importance of the intrinsic spectrin cytoskeleton for the integrity of GABA neurons in *C. elegans,* we used the auxin-inducible degron (AID) system (Ashley et al., 2021; L. Zhang et al., 2015) to degrade β-spectrin in a cell-specific and time-dependent manner. Degradation of β-spectrin from all neurons resulted in the widespread breakage of the DD and VD GABA neuron commissures. When the ability of axons to regenerate was inhibited, large portions of the dorsal nerve cord degenerated. By contrast, degradation of β-spectrin from the epidermis did not affect GABA neuron integrity. We also find that loss of β-spectrin from the epidermis does not affect the ability of axons to regenerate after laser axotomy. Together, these results demonstrate the cell-intrinsic importance of β-spectrin for axon integrity and establish targeted degradation of neuronal β-spectrin as a tool to study mechanisms of axon regeneration and degeneration.

## Results

### A system to acutely deplete β-spectrin and the spectrin-based membrane skeleton

To determine which tissues require β-spectrin to maintain GABA neuron integrity, we used targeted protein degradation (Ashley et al., 2021; L. Zhang et al., 2015) to specifically deplete β-spectrin from designated tissues in an otherwise wild-type background. We used an allele of *unc-70/*β-spectrin in which a small degron tag has been inserted into the endogenous *unc-70/*β-spectrin gene by CRISPR (UNC-70::AID) (Glomb et al., 2023) (Figure S1). We then expressed the E3 ligase TIR1 from *Arabidopsis thaliana* (atTIR1) under the pan-neuronal promoter *Prab-3*. Upon the addition of 1mM Auxin, atTIR1 will bind the degron tag resulting in rapid polyubiquitylation and proteasomal degradation (Holland et al., 2012; L. Zhang et al., 2015) of β-spectrin in all neurons. This allele has previously been used successfully to rapidly and reversibly degrade β-spectrin from the DA9 neuron (Glomb et al., 2023). Importantly, in the absence of auxin, the movement of the animals is normal, indicating that neither the degron tag on β-spectrin nor expression of atTIR1 perturb neuronal morphology or function (Figure 1A).

**Figure 1:**
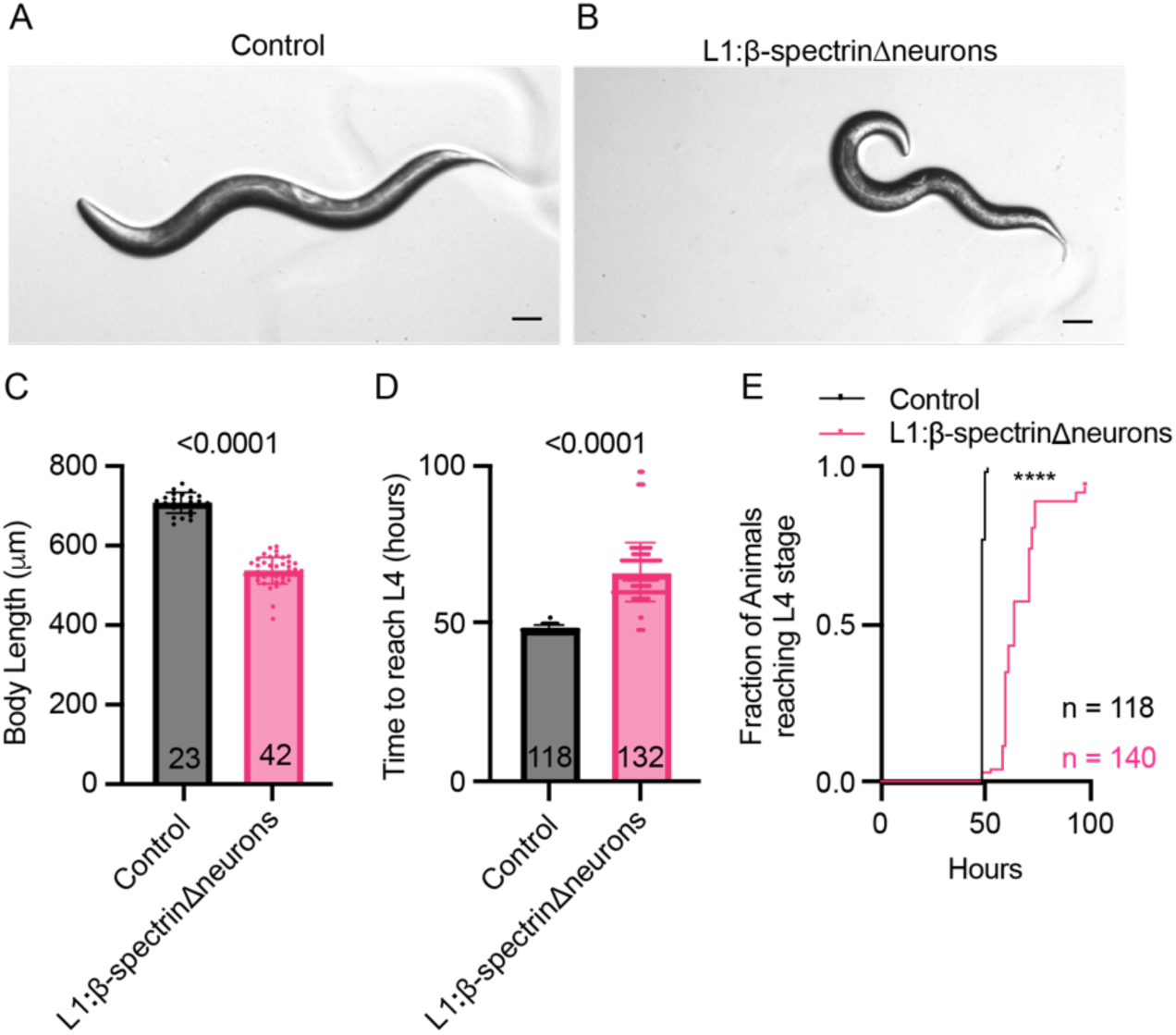
Degradation of neuronal β-spectrin from early larval stage causes morphological and developmental phenotypes. Representative images of L4 stage control animals homozygous for *unc-70::AID* and *P_rab-3_::atTIR1* and raised on (A) control or (B) 1mM auxin (IAA) to degrade β-spectrin from all neurons from the L1 stage (L1:β-spectrinΔneurons). Strain XE3230. Scale bar 50μm. (C) Quantification of the body length of control (mean = 704 μm) and L1:β-spectrinΔneurons animals (mean = 534 μm) at the L4 stage. (unpaired two-tailed t-test: p<0.0001) .(D) Quantification of the number of hours from egg lay to the L4 stage at 20°C in control (mean = 48.49 hours) and L1:β-spectrinΔneurons (mean = 65.95 hours) (unpaired two-tailed t-test: p<0.0001). Eight additional (5.7% of total) L1:β-spectrinΔneurons animals arrested at the L1/L2 stage and were censored at 98 hours. (E) Hours from egg lay to reach the L4 stage represented as a cumulative fraction of animals (Log-rank Mantel-Cox test: p<0.0001) The number of animals scored for each quantification is shown in or beside each dataset. Error bars show the standard deviation.

We allowed young adults to lay eggs on NGM plates containing 1mM natural auxin indole-3-acetic acid (IAA) for two hours. Newly hatched larvae are thus immediately exposed to auxin, allowing for pan-neuronal degradation of β-spectrin from the L1 larval stage (L1:β-spectrinΔneurons). (The eggshell has low permeability to IAA, therefore efficient degradation in the embryo is not expected (Negishi et al., 2019; L. Zhang et al., 2015)). We assessed these L1:β-spectrinΔneurons animals at the L4 stage and found that they exhibit abnormal body posture, reduced body length, and delayed development (Figure 1B-E).

To directly assess the effect of L1:β-spectrinΔneurons on the spectrin-based membrane skeleton (MPS), we imaged SPC-1/α-spectrin in GABA neurons, using an endogenously tagged split-GFP allele (Jia et al., 2019). In the MPS, α-spectrin and β-spectrin obligate dimers assemble into tetramers (Speicher & Marchesi, 1984; Yan et al., 1993). Expression levels of β-spectrin have been shown to affect expression of α-spectrin in *Drosophila* (Hülsmeier et al., 2007). Furthermore, it was previously shown that auxin-inducible degradation of β-spectrin results in eradication of the axonal MPS and loss of α-spectrin from axons of DA9 neurons (Glomb et al., 2023). Thus, axonal α-spectrin can be used as an endogenous marker for the presence of β-spectrin and an intact MPS.

We assessed L1:β-spectrinΔneurons animals 24 hours after egg lay, at which time both control and L1:β-spectrinΔneurons animals are still in the L1 larval stage. We found that α-spectrin was robustly localized in axons of GABA neurons in control animals (of the same genotype, but without auxin) (Figure 2B). (We would not expect to observe the periodic nature of the MPS under our imaging conditions due to resolution constraints.) However, in the presence of auxin, α-spectrin was absent from the axons of DD GABA neurons (Figure 2B,C). At this early larval timepoint, VD axon outgrowth had not yet occurred and therefore the MPS in VD axons could not be assessed. These data indicate that β-spectrin is rapidly degraded, disrupting the MPS and resulting in loss of α-spectrin from GABA neuron axons. Interestingly, we also observed accumulation of α-spectrin in the cell bodies of GABA neurons (Figure 2B, S2A,B) (see Discussion).

**Figure 2:**
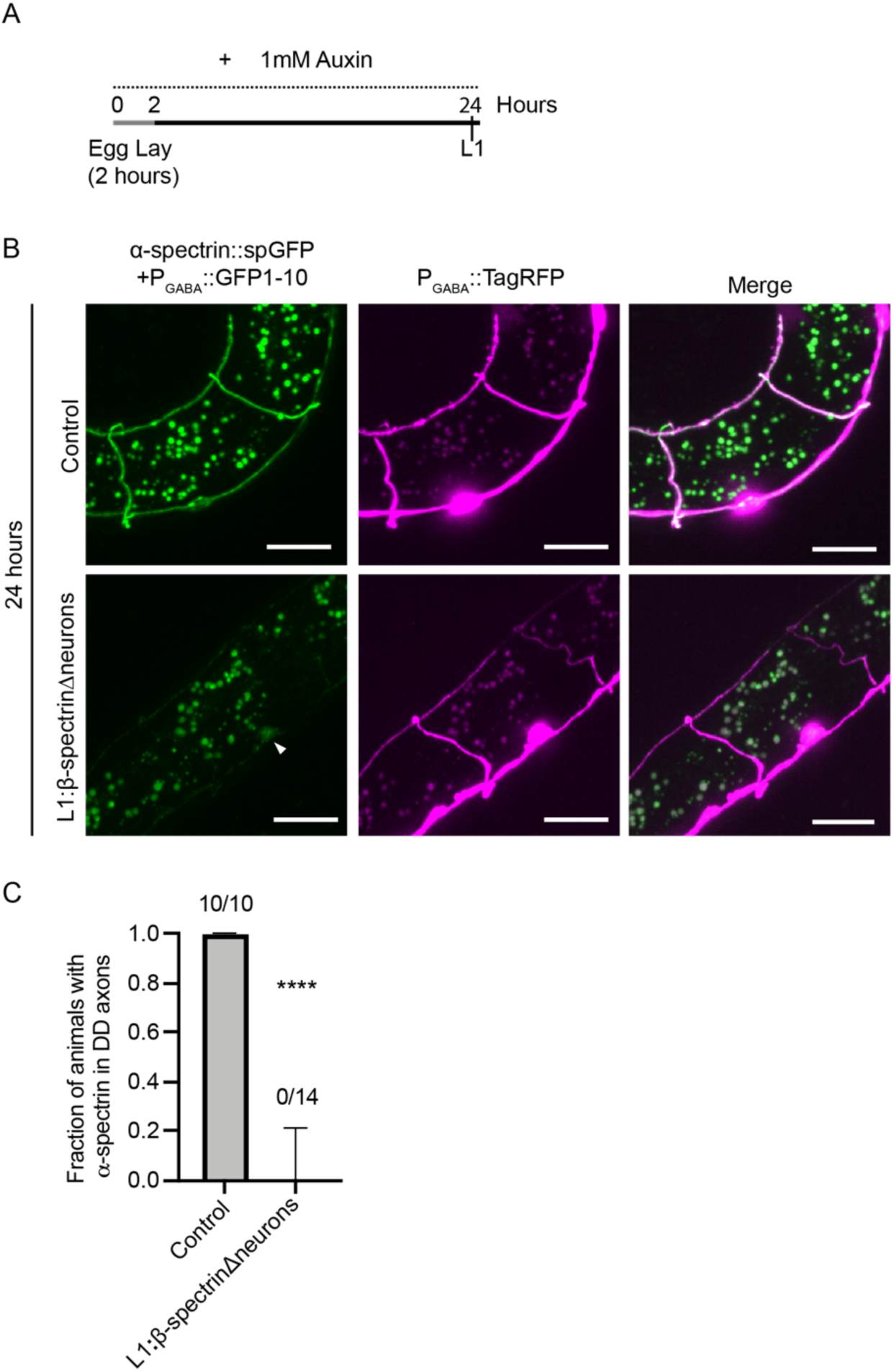
Degradation of neuronal β-spectrin disrupts the MPS within 24 hours. (A) Schematic of degradation timing for L1:β-spectrinΔneurons assayed at the L1 stage (B) Representative images of endogenous α-spectrin in GABA neurons (split-GFP labeled) and GABA neuron morphology (cytosolic TagRFP) in L1 control (no auxin) and L1:β-spectrinΔneurons animals (1mM auxin for 24 hours). Scale bar 10 μm. The white arrow points towards accumulated α-spectrin in cell body of L1:β-spectrinΔneurons animal. (C) Quantification of the fraction of animals with endogenous α−spectrin visible in any DD neuron commissure at the 24-hour time point (Fisher’s exact test: p<0.0001). The number of animals scored is shown above each bar.

### Axons break when neurons lack β-spectrin

To determine whether L1:β-spectrinΔneurons result in axon breaks, we visualized the VD/DD neurons at the L4 stage in animals raised on auxin. We found degradation of β-spectrin from all neurons resulted in widespread breakage of GABA neuron axons (Figure 3B-C). The axons of VD and DD-type GABA neurons typically overlap in the dorsal and ventral nerve cord, but their commissures are typically non-overlapping. Nearly 60% of axons were broken at larval stage L4, and 80% of animals exhibited gaps in the dorsal nerve cord (Figure 3B-D). Formation of the DD GABA neurons occurs embryonically before UNC-70 is degraded. By contrast, the VD GABA neurons are born post-embryonically and complete axon outgrowth by the L2 larval stage. To confirm that the phenotype is caused by axon breakage rather than axon outgrowth defect, we quantified the percentage of broken axons (Figure 3E) and number of visible intact VD/DD neuronal commissures (Figure 3F) at the L2 stage. At the L2 stage, when development is complete, nearly all commissures are intact (Figure 3E). By contrast, at L4 stage, most commissures are broken (Figure 3C). While there is a decrease in the number of intact commissures in L1:β-spectrinΔneurons at the L2 stage compared with control, this is likely due to spontaneous breakage shortly after or even during outgrowth (Figure 3F). This analysis reveals that loss of neuronal β-spectrin has minimal effect on outgrowth of the GABA neurons, but instead results in hyper-fragile axons that progressively break, consistent with previous results (Hammarlund et al., 2007).

**Figure 3:**
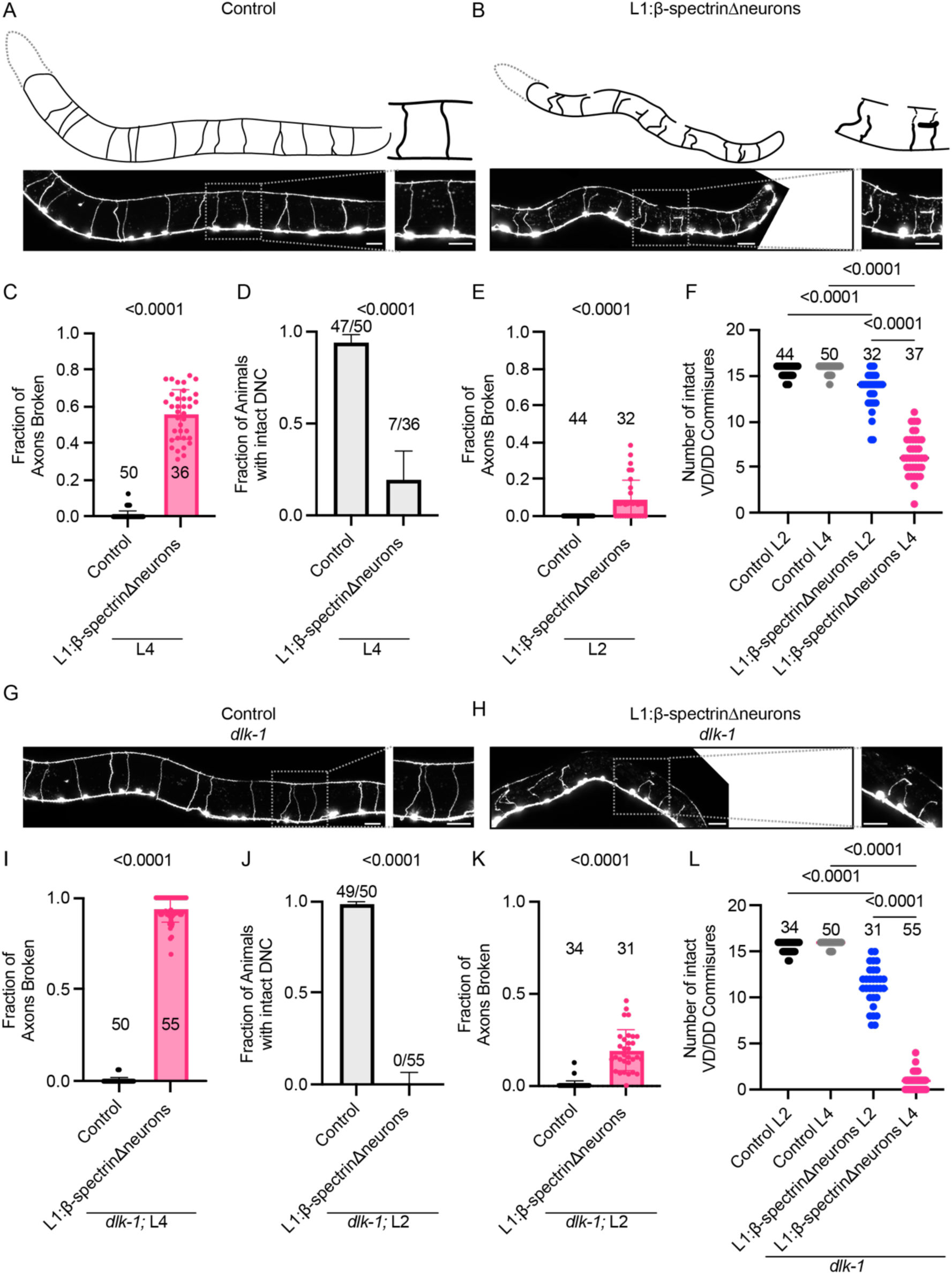
VD/DD axons break when neurons lack β-spectrin. (A-B) Representative images and cartoon tracings of VD/DD GABA neurons in (A) L4 stage control (no auxin) and (B) L4 animal raised on 1mM Auxin to degrade β-spectrin from all neurons from the L1 stage (L1:β-spectrinΔneurons). Strain XE3230. Scale bar 20μm. (C) Quantification of the fraction of axons broken per animal in L4 stage control and L1:β-spectrinΔneurons animals (unpaired two-tailed t-test: p<0.0001). (D) Quantification of the fraction of animals with an intact dorsal nerve cord (DNC) in L4 stage control and L1:β-spectrinΔneurons animals (Fisher’s exact test: p<0.0001). (E) Quantification of the fraction of axons broken per animal in L2 stage control and L1:β-spectrinΔneurons (unpaired two-tailed t-test: p<0.0001) (F) Quantification of the number of visible intact VD/DD commissures observed in L2 and L4 stage control and L1:β-spectrinΔneurons animals (One-way ANOVA followed by Tukey’s multiple comparisons test p<0.0001). (G-H) Representative image of VD/DD GABA neurons in L4 stage (G) *dlk-1* loss of function control (no auxin) and (H) *dlk-1* animal raised on 1mM Auxin to degraded β-spectrin from all neurons from the L1 stage (L1:β-spectrinΔneurons), Strain XE3303. Scale bar 20μm. (I) Quantification of the fraction of axons broken per animal in L4 stage *dlk-1* control and *dlk-1;* L1:β-spectrinΔneurons animals (unpaired two-tailed t-test: p<0.0001). (J) Quantification of the fraction of animals with an intact DNC in L4 stage *dlk-1* control and *dlk-1;* L1:β-spectrinΔneurons animals (Fisher’s exact test: p<0.0001) (K) Quantification of the fraction of axons broken per animal in L2 stage *dlk-1* control and *dlk-1;* L1:β-spectrinΔneurons animals (unpaired two-tailed t-test: p<0.0001). (L) Quantification of the number of intact VD/DD commissures observed in L2 and L4 control *dlk-1* and *dlk-1;* L1:β-spectrinΔneurons animals (ANOVA followed by Tukey’s multiple comparisons test: p<0.0001). The number of animals scored for each quantification is shown in or above each bar.

Degradation of neuronal β-spectrin resulted in robust regeneration of the GABA axons with many broken axons exhibiting growth cones (Figure 3B, S3B). This indicates that many axons that break also successfully regenerate (Hammarlund et al., 2007), complicating an accurate quantification of total axon breakage. To determine the full extent of axon breakage upon β-spectrin degradation, we introduced a mutation in the kinase *dlk-1* to prevent axon regeneration (Hammarlund et al., 2009). Loss of *dlk-1* together with L1:β-spectrinΔneurons revealed that 94% of axons break (Figure 3H, I) and 100% of animals have gaps in their dorsal nerve cord by the L4 stage (Figure 3J). In most animals, large portions of the dorsal nerve cord were fully degenerated (Figure 3H). Consistent with our observations for L1:β-spectrinΔneurons animals, *dlk-1;* L1:β-spectrinΔneurons animals were small (Figure 3H) and did not have large defects in GABA neuron outgrowth (Figure 3K,L). The phenotype of *dlk-1;* L1:β-spectrinΔneurons animals was consistent whether a genetic loss of function in *dlk-1* or a small molecular inhibitor of DLK-1 (GNE-3511) was used (Figure S3C,D). Thus, neuronal β-spectrin is essential for GABA neuron integrity, and loss of DLK-1 provides an improved way to quantify breakage.

### β-spectrin in the skin is not required for GABA neuron integrity or regeneration

Recently, epidermal β-spectrin has been proposed to protect touch receptor neurons (TRNs) and GABA neurons from breakage and degeneration (Coakley et al., 2022). Both TRNs and GABA motor neurons make close contacts with the epidermis. However, while the TRNs are embedded within the skin, the GABA neurons are not. To investigate the role of epidermal β-spectrin for GABA neuron integrity, we degraded β-spectrin from the epidermis using a variety of epidermal promoters (*P_col-10_:atTIR1* or *P_dpy-7_::atTIR1* and *P_col-19_:atTIR1*) (L1:β-spectrinΔskin) and confirmed degradation of epidermal β-spectrin was successful by observing disrupted epidermal α-spectrin (Figure S4). We observed no axonal breakage in L1:β-spectrinΔskin animals at the L4 stage (Figure 4A, B, D). To assess if axons had broken and regenerated, we used an inhibitor of DLK-1 (DLKi) to prevent regeneration. We observed only 2% of axons broken in L1:β-spectrinΔskin + DLKi animals (Figure 4B) in contrast to 98% in L1:β-spectrinΔneurons + DLKi animals at the L4 stage (Figure S3C). In a low proportion of animals we observed small gaps in the dorsal nerve cord when β-spectrin was degraded using the *col-10* promoter (24% of animals) (Figure 4A,C) or using the *dpy-7* and *col-19* promoters (8% of animals) (Figure 4E). This breakage was not exacerbated by inhibition of DLK-1 (Figure 4C). We hypothesize that these gaps are due to instability at the axon tips, rather than breakage along the axon shaft (see Discussion).

**Figure 4:**
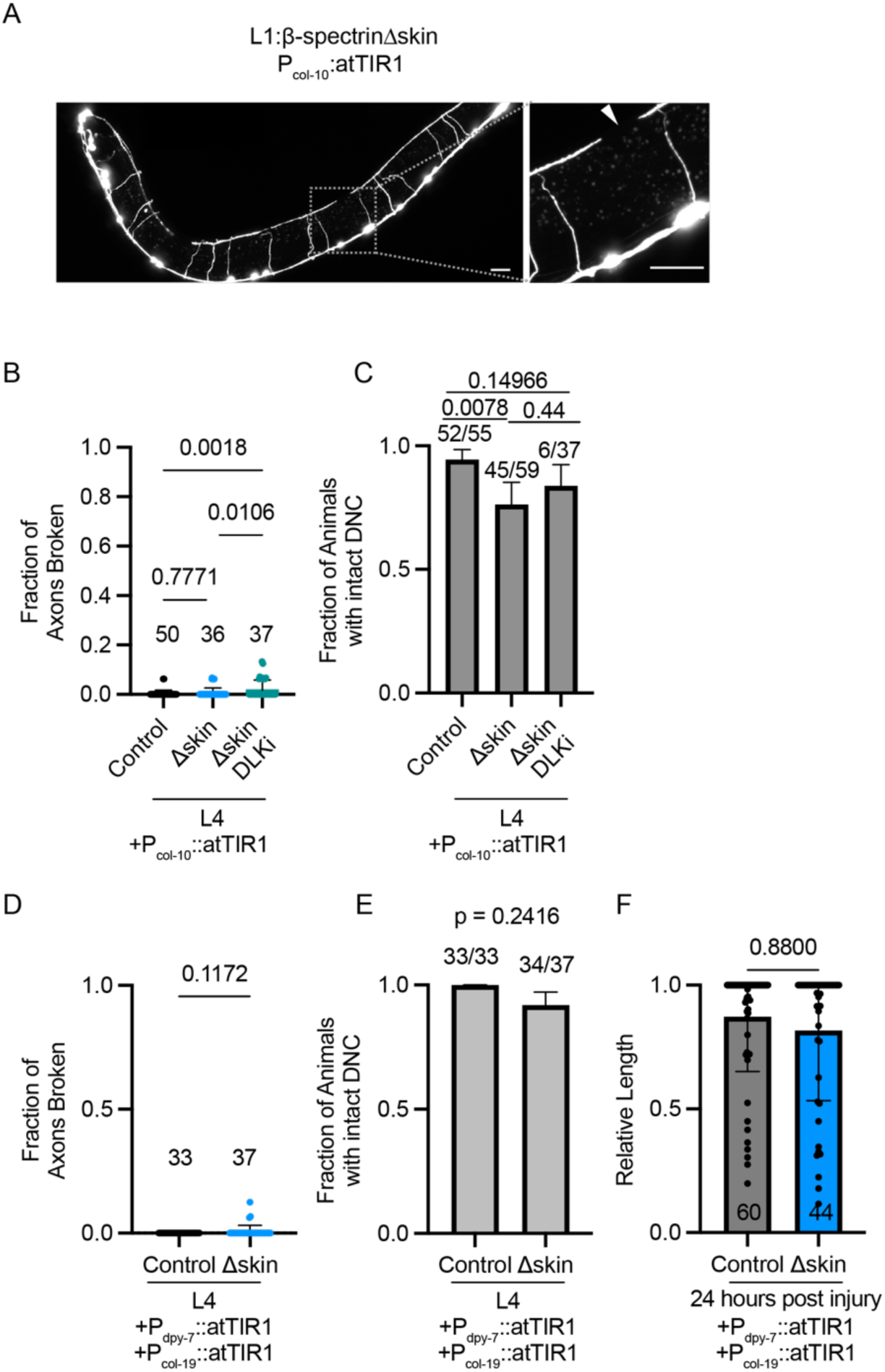
Epidermal β-spectrin is not required for GABA neuron integrity or regeneration. (A) Representative image of GABA neurons in L4 animal when β-spectrin is degraded from the epidermis (*P_col-10_::atTIR1* + 1mM Auxin) from the L1 stage (L1:β-spectrinΔskin). Scale bar 20μm. White arrow indicates gap in dorsal nerve cord. (B) Quantification of the fraction of axons broken when β-spectrin is degraded from the epidermis (L1:β-spectrinΔskin) using *P_col-10_::atTIR1* with and without inhibition of DLK (DLKi – 50μM GNE-3511). (ANOVA followed by Tukey’s multiple comparisons test). (C) Quantification of the fraction of animals with an intact dorsal nerve cord (DNC) in control, L1:β-spectrinΔskin, and L1:β-spectrinΔskin + DLKi using *P_col-10_::atTIR1* (Fisher’s Exact tests) (D) Quantification of the fraction of axons broken when β-spectrin is degraded from the epidermis (L1:β-spectrinΔskin) using *P_dpy-7_::atTIR1* and *P_col-19_::atTIR1* (unpaired two-tailed t-test: non-significant p = 0.1172). (E) Quantification of the fraction of animals with an intact dorsal nerve cord (DNC) in control and L1:β-spectrinΔskin using *P_dpy-7_::atTIR1* and *P_col-19_::atTIR1* (Fisher’s Exact test: p = 0.2416) (F) Quantification of axon regeneration as the relative length of regrowth 24 hours after laser-axotomy induced injury in control and Δskin animals (Kolmogorov-Smirnov test: nonsignificant p=0.88).

To investigate if epidermal spectrin plays additional roles in integrity or regrowth of motor neurons in the context of injury, we performed laser axotomy on the VD GABA neurons in L1:β-spectrinΔskin animals. Spectrin has been shown to regulate axon-glia attachments (Garcia-Fresco et al., 2006; Susuki et al., 2011, 2018) important for neuronal integrity, and β-spectrin regulates the attachment of touch receptor neurons to the epidermis (Coakley et al., 2020). The extracellular environment is known to affect the ability of VD/DD axons to regenerate after injury (Edwards & Hammarlund, 2014), and a previous report has visualized epidermal α-spectrin localized to regrowing PLM touch receptor neurons after injury (Coakley et al., 2022). Thus, it is possible that epidermal spectrin may influence VD axon regrowth after injury through either direct physical scaffolding or changes in the extracellular environment. However, L1:β-spectrinΔskin animals did not have a defect in the ability of axons to regrow after injury (Figure 4F). Together, these results demonstrate that epidermal β-spectrin does not have a large effect on GABA neuron integrity in basal or injury conditions.

### Cholinergic neurons, but not skin, modulate the effect of GABA neuron spectrin loss

We next sought to determine if loss of β-spectrin only from GABA neurons caused similar breakage to pan-neuronal degradation. We used the GABA-specific *unc-47* promoter to drive expression of atTIR1, and the *dlk-1* mutant background to block regeneration and allow accurate quantitation. Interestingly, loss of β-spectrin only from GABA neurons (L1:β-spectrinΔGABA) resulted in 100% of animals with degeneration in their dorsal nerve cord (Figure 5A,C) but only approximately 50% of axons broken at the L4 stage (Figure 5B). Multiple atTIR1 arrays gave similar phenotypes (Figure S5A), and α-spectrin distribution was disrupted in all GABA commissures regardless of axon breakage (Figure S5B,C). Thus, loss of β-spectrin from GABA neurons is sufficient to cause axon breakage, but breakage is enhanced when β-spectrin is lost from all neurons.

**Figure 5:**
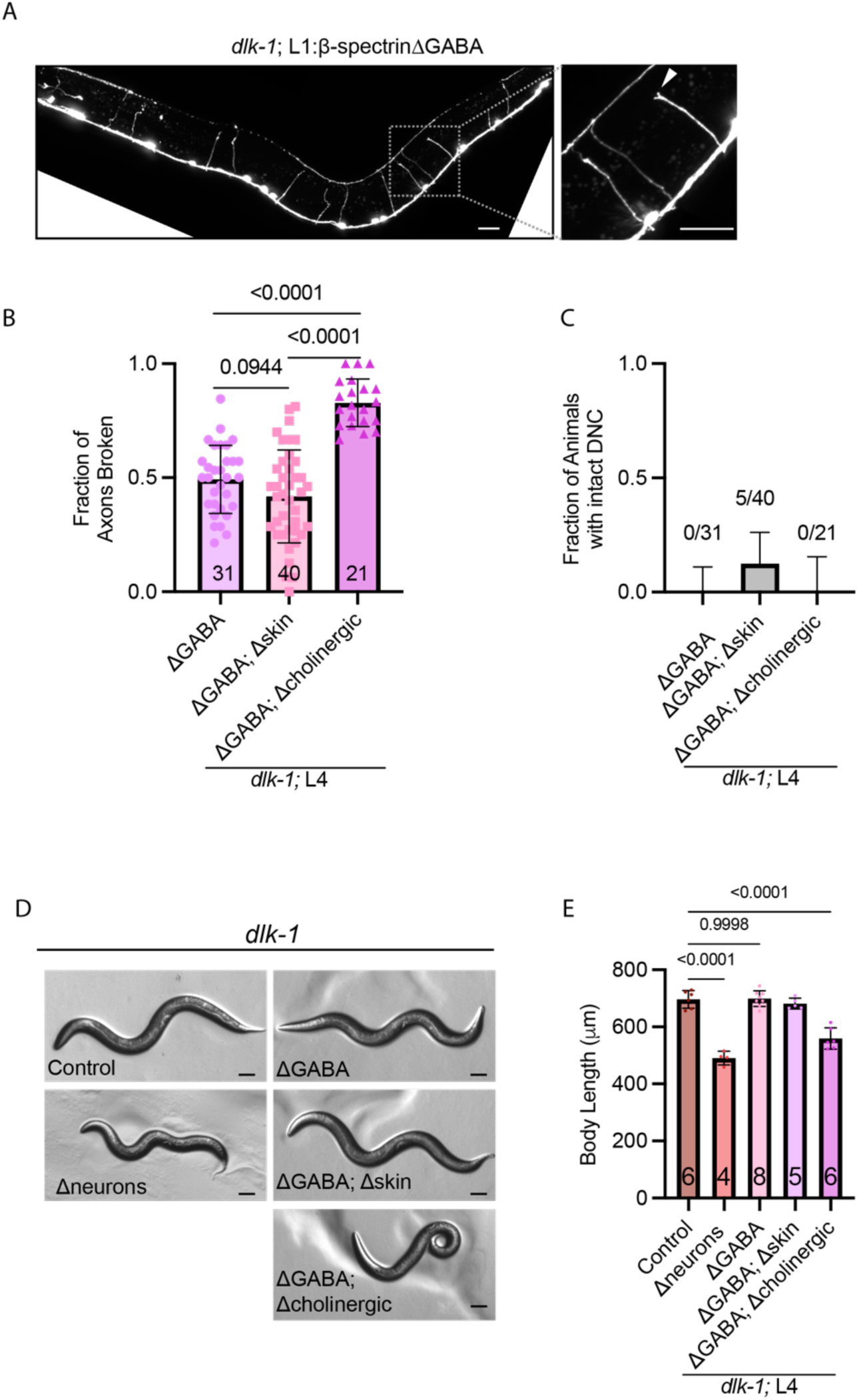
β-spectrin in cholinergic neurons modulates the effect of GABA neuron spectrin loss on axon integrity. (A) Representative image of *dlk-1* animal raised on 1mM Auxin to degraded β-spectrin from GABA neurons only (L1:β-spectrinΔGABA). Scale bar 20μm. White arrow indicates broken axon. (B) Quantification of the fraction of axons broken per animal in L4 *dlk-1* animals when β-spectrin is degraded from GABA neurons only (*P_unc-47_::atTIR1*), GABA neurons and the skin (*P_unc-47_::atTIR1*, *P_dpy-7_::atTIR1* and *P_col-19_::atTIR*), and GABA neurons and cholinergic neurons (*P_unc-47_::atTIR1*, *P_unc-17_::atTIR1*) (ANOVA followed Tukey’s multiple comparisons test: p<0.0001) (C) Quantification of the fraction of animals with an intact dorsal nerve cord (DNC) when β-spectrin is degraded from GABA neurons only, GABA neurons and the skin, and GABA neurons and cholinergic neurons (D) Representative images of body morphology and (E) quantification of body length in L4 stage animals when β-spectrin is degraded from all neurons, GABA neurons, GABA neurons and skin, and GABA neurons and cholinergic neurons in the *dlk-1* loss of function background. Scale bar 50μm. Control animals contain the *P_rab-3_::atTIR1* allele but were not exposed to auxin. (ANOVA followed Tukey’s multiple comparisons test: p<0.0001).

To further explore how loss of GABA spectrin might be enhanced by loss of spectrin from other tissues, we degraded β-spectrin from GABA neurons in combination with two other tissues: the skin or the cholinergic neurons. Degradation of β-spectrin from both GABA neurons and the skin did not exacerbate breakage when compared to GABA alone (Figure 5B). However, degradation from both GABA and cholinergic neurons resulted in 83% of axons broken, similar to that observed with pan-neuronal degradation (Figure 5B).

How might loss of β-spectrin from cholinergic neurons enhance axon breakage in GABA neurons that also lack β-spectrin? We considered the possibility that the cholinergic neurons might physically scaffold the GABA neurons. Cholinergic motor neurons such as VA and VB also extend commissures across the body of the worm, and these axons can fasciculate with GABA neuron axons. In this model, the elimination of β-spectrin only in GABA neurons might have a weaker effect because undamaged cholinergic neurons are available as scaffolds. To test this model, we visualized the cholinergic neuron commissures in L1:β-spectrinΔGABA animals using the pan-cholinergic marker P_unc-17_::TagRFP or P_unc-17_::mTagBFP2 (Figure S5D,F). We found that fasciculation events are rare, with only 1-2 of the 19 GABA commissures in each animal fasciculated with a cholinergic commissure. Thus, the scaffolding model cannot explain the approximately 8 unbroken commissures per animal when spectrin is lost from only GABA neurons. While we observed the majority of fasciculated GABA commissures are intact, which suggests a possible correlation between fasciculation and preventing breakage (Figure S5D,E), we also observed fasciculated GABA commissures that had broken (Figure S5E, F). Therefore, fasciculation is not sufficient to prevent breakage of GABA neurons caused by degradation of β-spectrin. Together, these results indicate that scaffolding by fasciculation may inhibit axon breakage but is not the primary way that β-spectrin in cholinergic neurons acts to limit breakage in GABA neurons.

Next, we considered the possibility that differences in movement might underlie the role of cholinergic β-spectrin in GABA neuron breakage. Degradation of β-spectrin from only GABA neurons resulted in a mild movement phenotype (data not shown) and no significant effect on body size (Figure 5D,E). In contrast, degradation of β-spectrin from both GABA and cholinergic neurons reduced body size and strongly affected movement (Figure 5D,E). These animals are hyper-curled, similar to β-spectrinΔneurons animals (Figure 5D). We hypothesize the increased breakage in β-spectrinΔneurons and β-spectrinΔGABAΔcholinergic animals is due to increased physical strain on the GABA neurons resulting from altered body movement and posture. This is consistent with previous results showing that the strain of movement induces breakage in hyper-fragile axons (Hammarlund et al., 2007).

### Spectrin is required in mature neurons to prevent axon breaks

Although there have been multiple reports of axon damage due to loss of β-spectrin (Coakley et al., 2020, 2022; Hammarlund et al., 2007; Krieg et al., 2014, 2017), in general, these studies relied on constitutive loss of β-spectrin. In the above experiments, we exposed animals to auxin at the L1 stage, triggering β-spectrin degradation while the nervous system is still developing. In particular, the VD GABA neurons are still extending growth cones at this stage, while the DD GABA neurons still must undergo extensive synaptic remodeling (Hallam & Jin, 1998; White et al., 1978). We next examined whether acute loss of β-spectrin in mature neurons is sufficient to cause axon breakage. We exposed β-spectrinΔneurons animals to auxin at the L4 stage, at which time the GABA neurons are fully mature. After 48 hours, approximately 50% of VD/DD axons were broken, with the majority exhibiting evidence of regrowth, including large growth cones (Figure 6A, C). Thus, like developing neurons, mature neurons depend on intrinsic neuronal β-spectrin to preserve axon integrity.

**Figure 6:**
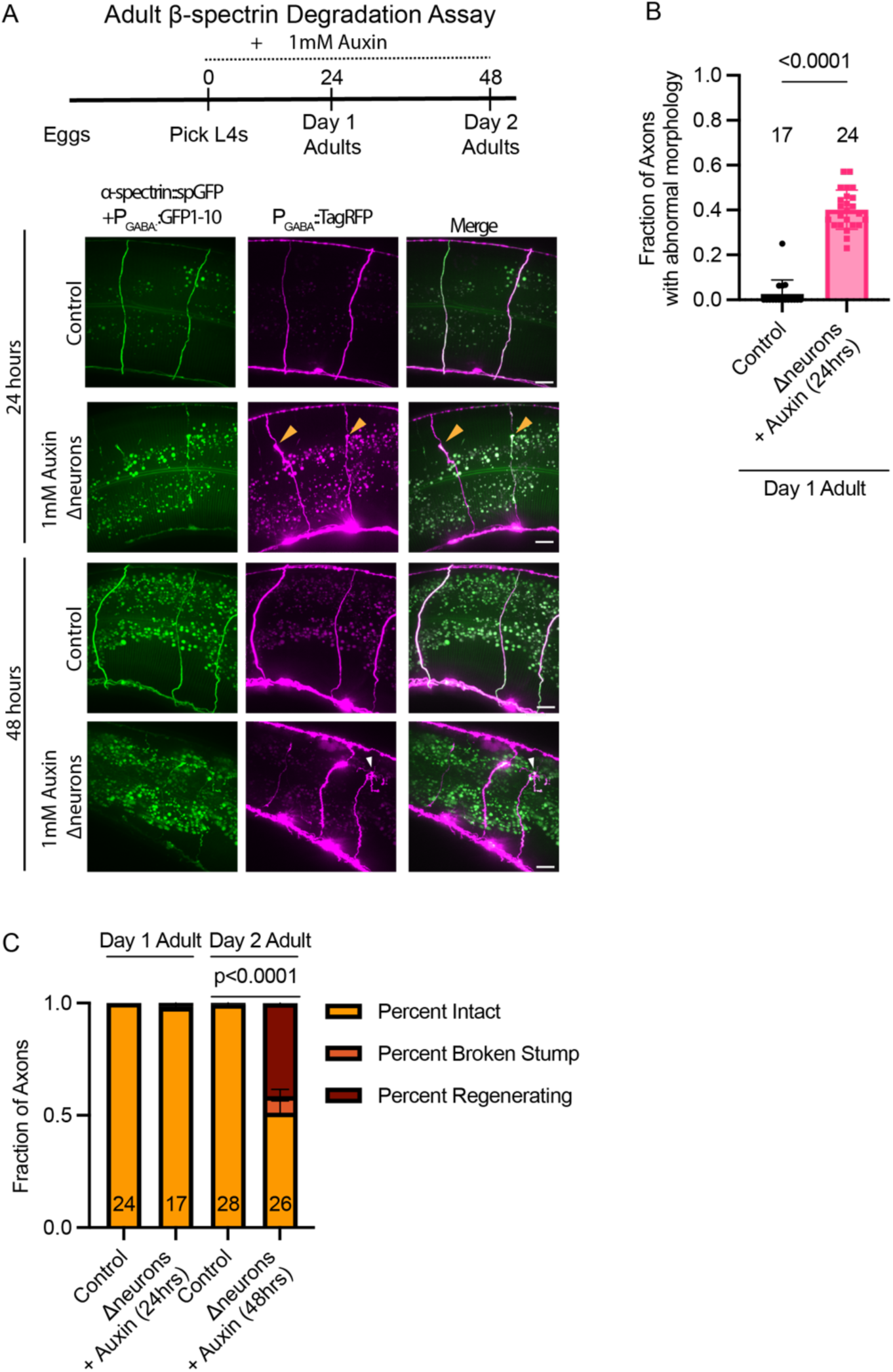
Spectrin is required in mature neurons to prevent axon breaks. (A) Adult β-spectrin degradation assay: Representative images of endogenous α-spectrin in GABA neurons (split-GFP) and GABA neuron morphology (cytosolic TagRFP) in day 1 and day 2 control and β-spectrinΔneurons adults when β-spectrin is degraded starting at the L4 stage (time 0) Scale bar 10 μm. Yellow arrows indicate intact axons with abnormal morphology (axon thinning). White arrow indicates a broken and regenerating axon. (B) Quantification of the fraction of axons with abnormal morphology after 24 hours of β-spectrin degradation from L4 stage to day 1 of adulthood. (unpaired two-tailed t-test: p<0.0001). The number of animals scored is shown above each bar. (C) Quantification of the fraction of axons intact, broken and regenerating, broken and non-regenerating (stump) when β-spectrin is degraded for 24 or 48 hours from L4 stage to day 1 and 2 of adulthood. (Fisher’s exact test: p<0.0001). The number of animals scored is shown inside each bar.

### Axon breakage is delayed following neuronal spectrin disruption

Finally, we investigated the temporal relationship between loss of the MPS and axon breakage. We degraded β-spectrin for 24 or 48 hours in both the developing and mature nervous system. In both developing and mature neurons, α-spectrin distribution was completely disrupted within 24 hours of auxin treatment, indicating that AID activation rapidly degrades β-spectrin and disrupts the MPS (Figure 2, Figure 6). Strikingly, however, breakage was not observed at 24 hours, but was robust at 48 hours (Figure S2C, S3C, 6C). For example, in L1:β-spectrinΔneurons animals 24 hours after egg lay onto auxin plates, α-spectrin distribution was disrupted (Figure 2); however, we found no visible breakage of the DD neurons (at this timepoint the VD axons had not yet developed) (Figure S2C). After 48 hours of auxin treatment, L1:β-spectrinΔneurons animals exhibit widespread breakage (Figure 3, S3). Similarly, in the mature nervous system, L4 animals exposed to auxin for 24 hours had significantly disrupted α-spectrin (Figure 6A, S6A), and while axon morphology was abnormal including large swellings and thinnings (Figure 6A,B), axons were continuous and not visibly broken. By 48 hours, approximately half of all axons were visibly broken (Figure 6A,C). As in the larval stage animals, we also observed α-spectrin accumulation in the cell bodies when β-spectrin is degraded (Figure S6B,C). Therefore, in both the developing and mature nervous systems, MPS disruption significantly precedes axon breakage.

## Discussion

### The neuron-intrinsic spectrin cytoskeleton is required for motor neuron axon integrity during typical and atypical movement

We have shown that the intrinsic MPS is required for VD/DD GABAergic motor neuron integrity in *C. elegans*. Degradation of β-spectrin during development and in the mature nervous system altered α-spectrin distribution and resulted in breakage of VD/DD commissures. We observed α-spectrin accumulated in the cell bodies when β-spectrin was degraded. This is consistent with previous observations in DA9 (Glomb et al., 2023). Without β-spectrin, α-spectrin fails to be properly localized in the axon, remains highly mobile (Glomb et al., 2023), and the MPS cannot be formed. Interestingly, degradation of β-spectrin only from GABA neurons resulted in less breakage than degradation from all neurons or degradation from both GABA and cholinergic neurons. We hypothesize this is due to more stressful body posture and movement when β-spectrin is degraded from cholinergic neurons. Previous work has shown that the DVA neuron is activated during compressive stress caused by body bending, and cell intrinsic loss of *unc-70* using the Cre-lox system from the DVA neuron alone caused an “exaggerated” body posture characterized by deeper body bends (Das et al., 2021). Similar body posture was observed in both β-spectrinΔneurons and β-spectrinΔGABAΔcholinergic animals (Figure 5D). These deep body bends may place additional stress on VD/DD axons, particularly where the commissures squeeze between the muscle and the skin (Knobel et al., 1999), resulting in increased frequency of breakage in neurons that lack β-spectrin. Overall, our data demonstrate that intrinsic β-spectrin is required during both normal movement and stressful movement to maintain motor neuron axon integrity.

### Neuronal rather than epidermal spectrin protects GABA neurons from breakage

The importance of the intrinsic MPS has been shown across model organisms. Loss of αII-spectrin from CNS neurons in mice causes widespread neurodegeneration and seizures (Huang, Zhang, Ho, et al., 2017), and mice lacking αII-spectrin in the PNS have disrupted nodes of Ranvier, mislocalized Kv1 K+ channels, degeneration of large-diameter myelinated axons, and severe ataxia (Huang, Zhang, Zollinger, et al., 2017). Mathematical modeling describes how spectrin tetramers within the MPS may provide mechanical neuroprotection through reversibly unfolding to act as “shock absorbers” and buffer tension (Dubey et al., 2020). In *C. elegans*, the MPS is under continuous tension in living neurons, and if this tension is disrupted by mutations in β-spectrin, touch sensation is impaired (Krieg et al., 2014). Neurons in culture withstand compression, torsion, and stretch through tension provided by the MPS and stiffness provided by the microtubule cytoskeleton (Krieg et al., 2017). However, the importance of the intrinsic MPS for axon integrity in *C. elegans* may vary by neuron type, morphology, and environment. Loss of β-spectrin from DVA only causes axon buckling, not breakage or degeneration (Das et al., 2021). Touch receptor neurons buckle in β-spectrin mutants, and while expression of epidermal β-spectrin can decrease buckling, it does not rescue touch response, supporting the importance of the intrinsic MPS for neuronal function (Krieg et al., 2014).

The *C. elegans* system offers numerous advantages for studying the importance of both the intrinsic MPS and spectrin in surrounding tissues for neuronal function and integrity in a free-living intact organism. Both TRNs (including the PLM neuron) and VD/DD GABAergic motor neurons are in contact with the epidermis. The epidermis serves a glial-like function including clearance of axonal fragments after injury and mediating communication with the extracellular environment (reviewed in (Chisholm & Xu, 2012)). A spectrin scaffold within the epidermis has also been shown to protect the integrity of TRNs, which are embedded in the epidermis and mediate response to touch (Bonacossa-Pereira et al., 2022; Coakley et al., 2020, 2022). While VD/DD axons are not embedded in the epidermis, they are in close contact with both the epidermis and muscle. In a hypergravity-induced model of axon damage, epidermal and muscle β-spectrin and the hemidesmosomes connecting axons to the epidermis promote GABAergic motor neuron axon defects that occur most frequently along larger muscle cells (Kalichamy et al., 2020). There it is possible that scaffolding and attachment to the epidermis may support VD/DD axon integrity.

To test the contribution of epidermal β-spectrin for GABA neuron integrity, we degraded epidermal β-spectrin using a variety of epidermal promoters (P_col-10_ or P_dpy-7_ and P_col-19_:), none of which resulted in significant axon breakage (Figure 4). Additionally, degradation of epidermal β-spectrin did not affect the guidance or regrowth of axons after injury (Figure 4F). We did observe a small fraction of L1:β-spectrinΔskin animals with small gaps in the dorsal nerve cord (Figure 4A,C,E). However, we hypothesize these gaps are due to instability of the axon tips, not breakage. Gaps were not exacerbated by inhibition of regeneration (Figure 4C) as they are in L1:β-spectrinΔneuron animals (Figure S3D). The DD and VD neurons each separately tile the dorsal nerve cord, with the tips of adjoining processes connected by gap junctions. Epidermal β-spectrin may function to promote or stabilize the establishment of this tiling pattern.

Recent results have suggested that β-spectrin in the epidermis is required to protect against breakage in GABA neurons (Coakley et al., 2022). That study used the FLP-FRT system, activated by the *dpy-7* epidermal promoter, to delete the *unc-70* gene in skin cells. However, in addition to terminally-differentiated epidermal cells, *dpy-7* is also expressed earlier in the P cell lineage, from which both epidermal cells and the VD GABA neurons are derived (Gilleard et al., 1997; Sulston & Horvitz, 1977). Since FLP acts at the DNA level, when a precursor cell is edited, all cells derived from it will also contain the edit. As a consequence, expression of FLP under the *dpy-7* promoter would be expected to delete β-spectrin in both VD GABA neurons and in epidermis. Thus, we hypothesize that in the previous experiments, as in ours, removal of β-spectrin from GABA neurons results in axon breaks. Consistent with this result, previous work showed rescue of β-spectrin in the epidermis did not rescue VD/DD neuron breakage in a loss-of-function β-spectrin mutant (Hammarlund et al., 2007). Additional experiments will need to be done to clarify the differences between these two methods of cell-specific targeted loss of β-spectrin. Nevertheless, our results confirm the intrinsic importance of the MPS for VD/DD GABAergic motor neuron integrity throughout lifespan.

### Loss of the GABAergic MPS as a model to study axon regeneration and degeneration *in vivo*

We show here that loss of β-spectrin from all neurons causes complete breakage of VD/DD GABAergic neurons, and that loss of β-spectrin from only GABA neurons also results in significant (though incomplete) breakage of VD/DD GABAergic neurons. In both cases, the animals are much healthier than full body β-spectrin mutants (Hammarlund et al., 2000) which have previously been used successfully in unbiased screens to identify conserved mediators of axon regeneration (Hammarlund et al., 2009; Nix et al., 2014). Thus, these animals may enable new approaches to studying axon regeneration and degeneration, in which targeted depletion of neuronal β-spectrin is performed during development or in the mature nervous system to allow for high throughput screening of genes or compounds that affect axon regeneration and degeneration. In addition, the AID system may allow β-spectrin to be acutely degraded and then the animals removed from auxin, preserving the stability of newly-synthesized β-spectrin and allowing the study of MPS formation *in vivo*.

While previous studies have highlighted the importance of spectrin and the MPS for axon regeneration (Hofmann et al., 2022), it is not possible to carefully discriminate if intrinsic β-spectrin or the MPS promotes regeneration in our system because without β-spectrin axons break; however, the robust growth cone formation in L1:β-spectrinΔneurons and L1:β-spectrinΔGABA animals demonstrates intrinsic β-spectrin is not required for growth cone initiation or outgrowth, consistent with previous data (Hammarlund et al., 2007, 2009; Nix et al., 2014). In fact, it is possible that loss of β-spectrin may enhance axon regeneration. Disruption to the cytoskeleton through destabilizing actin or microtubules has been shown to activate the DLK/JNK pathway and mimic a pre-conditioning injury and promote axon regeneration (Valakh et al., 2015). Thus, the robust regeneration we observe in neurons that lack β-spectrin may be due to hyperactivation of DLK/JNK in L1:β-spectrinΔneurons animals.

Surprisingly, we observed that the MPS is disassembled hours before breakage and degeneration occur. These data suggest that the MPS is not the only structure that promotes neuronal integrity. We hypothesize that in the absence of the MPS, mechanical strain causes damage to neurons to accumulate over time, eventually resulting in axonal breaks. Identifying the processes downstream of MPS loss that control axon breakage would help elucidate how axons act to preserve their integrity.

## Methods

### Caenorhabditis elegans

*C. elegans* were maintained on standard nematode growth media (NGM) seeded with *Escherichia coli* OP50 at 20°C unless otherwise stated. Hermaphrodites were used for all experiments. Males were used only for crosses. Strains used in this work are listed in Table S1.

### Generation of transgenic animals

Transgenic animals were generated by microinjection (Stinchcomb et al., 1985). Promega 1kb DNA ladder was used as filler in the injection mix. In most cases*, P_myo-2_::mCherry* (1.0 ng/uL) was used as a co-injection marker. Plasmids used to generate transgenics are listed in Table S2. Multicopy targeted insertion to generate the *wpIs495*[*P_rab-3_::atTIR1::let-858UTR + P_myo-2_::GFP*] allele was performed as previously described (El Mouridi et al., 2022). Briefly, we performed CRISPR-Cas9 mediated targeted extrachromosomal array integration into the modular safe-harbor transgene insertion landing site (moSCI) locus on chromosome IV in strain CFJ94. Successful integrates were screened for rescue of *unc-119* movement phenotype and insertion was verified by PCR (El Mouridi et al., 2022).

### Molecular Cloning

Plasmids used for expression of transgenes in *C. elegans* were assembled into the destination vector pDEST^TM^R4-R3 (Invitrogen) using Gateway recombination (Invitrogen). Entry clones for Gateway LR reactions were generated by Gibson Assembly (Gibson et al., 2009). All plasmids used in this work are listed in Table S2.

### Auxin-inducible degradation

All auxin-induced knock-down experiments were performed by culturing animals on NGM plates containing 1 mM Auxin (#A10556.14; Thermo Fisher Scientific) seeded with OP50. Unless otherwise stated, gravid young adult animals were placed on 1mM Auxin plates and progeny were allowed to develop at 20°C. Animals of the same stage cultured on NGM plates lacking auxin were controls. Because animals raised on 1mM Auxin to degrade pan-neuronal β-spectrin are significantly smaller than control animals, visualization of vulva development was used to confirm analysis was performed at the L4.4 – L4.7 stage unless otherwise stated (Mok et al., 2015). For timed assays, one-day-old gravid adults were allowed to lay eggs on 1mM auxin-containing plates or control plates for two hours. Adults were subsequently removed, and progeny were allowed to develop at 20°C. Time 0 is defined as the start of the two-hour egg lay. Progeny were evaluated at the late L1 stage, exactly 24 hours after the start of the egg lay. To degrade β-spectrin in the mature nervous system, L4 stage animals were placed on 1mM Auxin plates (time 0) and analyzed 24 hours (day 1 adult) or 48 hours (day 2 adult) later. L4 animals of the same genotype placed on control NGM plates (time 0) served as controls.

### Fluorescence microscopy

Animals were immobilized using 10mM Levamisole (Santa Cruz, sc-205730) and mounted on a 3% (wt/vol) agarose pad on a glass slide. Confocal imaging was performed using a laser Safe DMi8 inverted microscope (Leica) equipped with a VT-iSIM system (BioVision), an ORCA-Flash 4.0 camera (Hamamatsu), HC PL APO 63x/1.40NA OIL CS2, a HC PL APO 40x/1.30NA OIL CS2, a HC PL APO 20x/0.8NA AIR and HC PL APO 100x/ 1.47NA OIL objective as well as 488 nm, 561 nm and 637 nm laser lines. Image acquisition was controlled by MetaMorph Advanced Confocal Acquisition Software Package. A step size of 0.2 μm was used. In cases where only axon morphology or breakage was being assessed, we used an Olympus BX61 microscope with attached Olympus DSU, Olympus UPlanFL N 40X/1.30 oil objective or UPlanSApo 60X/1.35 oil objective, and Hamamatsu C11440 camera. Image acquisition was controlled by Micro-Manager with a 0.4 μm step size.

### Analysis of fluorescence microscopy

Raw images were processed and analyzed with Fiji/ImageJ v2.16.0/1.54g. Images were acquired as a single image or multiple z layers and stacked into maximum projections using the maximum intensity. To visualize the entire axon or animal that could not be captured in a single field of view, multiple images were acquired and stitched together into a single image using the pairwise-stitching plugin with a linear blending fusion method (Preibisch et al., 2009). Analysis was performed on L4 stage animals unless otherwise noted.

### Quantification of axon breakage and gaps

Axon breakage was identified as a discontinuity of the cytosolic fluorescent marker (GFP, TagRFP, or chFP). Axons that were thin, dim, swollen, or beading were quantified as intact. If these abnormal morphologies were present, they were quantified separately from breaks/gaps and discussed as such in the results.

### Laser axotomy and quantification of axon regeneration

Laser axotomy was performed as previously described (Byrne et al., 2011, 2014; Kosmaczewski et al., 2015). Briefly, worms of the same genotype (strain XE3449) were allowed to lay eggs on plates with 1mM auxin (L1:β-spectrinΔskin) and without (control). Progeny were allowed to develop and larval stage L4.4-L4.6 animals carrying the wpEx570 array were mounted on a 3% (wt/vol in M9 buffer) agarose pad, immobilized by 0.05 mm microbeads (Polysciences, #08691) with 0.02% SDS, and visualized with a Nikon Eclipse 80i Microscope with a 100x Plan Apo VC lens (1.4 NA). Fluorescently-labeled ventral type (VD) GABA motor neuron commissures were severed at the dorsoventral midline using a 435 nm Photonic Instruments Micropoint laser with 10 pulses at 20 Hz. Three to four of the seven most posterior commissures were severed per worm. Worms were recovered at 20°C to NGM plates seeded with OP50 with 1mM auxin (L1:β-spectrinΔskin) or without (control) and scored for axon regeneration 24 hours after axotomy.

To score regeneration, worms imaged by confocal microscopy. Images were blinded to treatment group prior to analysis. Axon regeneration was scored as the relative length of regrowth. This is defined as the vertical distance between the ventral nerve cord and the growth cone or stump of the injured axon, normalized to the corresponding vertical distance between the dorsal and ventral cords as previously described (Zeng et al., 2018). The relative lengths were measured with ImageJ and plotted with GraphPad Prism. Full regeneration to the dorsal nerve cord corresponds to a relative length value of 1. If an axon appeared to fuse with the distal stump, it was given a score of full regeneration (relative length = 1). Because the data are not normally distributed, the non-parametric Kolmogorov-Smirnov test was used to compare the cumulative distribution of the normalized axon regrowth measurements between control and experimental groups.

### Measurement of animal length and body morphology

L4 animals were picked onto unseeded NGM plates and the plate was placed under a Leica M165FC stereoscope. Still images of individual animals were captured with a Basler acA2440 camera controlled by the WormLab software. Images were analyzed in ImageJ and the length of each worm was measured from the base of the tail to the mouth.

### Timed development assay

Thirty Day 1 gravid adults were allowed to lay eggs for 2 hours on NGM plates with and without 1mM auxin. All adults were removed at the end of the 2 hours. Progeny were allowed to develop at 20°C. The number of progeny reaching the L4 stage was quantified every two hours from 44 hours 64 hours, every two hours from 70 hours to 82 hours, and again at 94 hours and 98 hours. Once a worm was counted as reaching the L4 stage it was removed from the plate to avoid double counting. The average time of each animal to reach the L4 stage was quantified. The proportion of animals reaching the L4 stage at each timepoint was graphed using GraphPad Prism and compared using a log-rank (Mantel-Cox test). Any animal developmentally arrested and not reaching the L4 stage by 98 hours was censored.

### Inhibition of DLK using a small molecular inhibitor

DLK-1 was inhibited using the GNE-3511 inhibitor (MedChemExpress, HY-12947). The inhibitor was dissolved in DMSO and added to 50°C NGM media at a final concentration of 50μM prior to pouring. 1mM Auxin was also added to the media prior to pouring. Animals were allowed to lay eggs on plates containing 1mM Auxin + 50μM GNE-3511. Progeny were allowed to develop and were analyzed at the L4 stage.

### Statistical analysis

All statistical analysis was performed in GraphPad Prism. For non-binary data, unless otherwise noted, two-sided *t* test was used to compare two groups, and one-way ANOVA test followed by Tukey’s multiple comparisons test was used to compare multiple groups. Bar graphs depict mean, and error bars show standard deviation. The number of animals scored for each quantification is shown in or beside each dataset. Unless otherwise stated, data distributions were assumed to be normal, but this was not formally tested. For relative length quantification of regeneration following laser axotomy, data was not normally distributed; therefore, the non-parametric Kolmogorov-Smirnov test was performed to compare cumulative distributions. For binary data, Fisher’s exact tests were performed.

### Materials availability

Strains and plasmids used in this study are listed in Tables S1 and S2 respectively and will be made available to the scientific community upon request directed to Dr. Hammarlund. Strains generated in this study will be deposited at the Caenorhabditis Genetics Center (CGC).

### Data availability

All data are available in the main text or the supplementary materials.

## Acknowledgements

We thank all members of the Hammarlund and Yogev Labs for discussions and technical advice. We thank Massimo Hilliard and Sean Coakley for sharing the *vdSi17[Punc-25::ChFP] IV* allele used in strains XE3465 and XE3449.

This research was supported by the Gruber Science Fellowship at Yale and NINDS F31NS127495 to CAD, and by NINDS 5R01NS094219 to MH. Some strains were provided by the CGC, which is funded by NIH Office of Research Infrastructure Programs P40OD010440. WormBase was used in the planning, design, and analysis of this research (WS295, 2024) (Sternberg et al., 2024) and is supported by NHGRI U24HG002223. Gene expression and splicing information used in the design and analysis of this work was provided by The Complete Gene Expression Map of the *C. elegans* Nervous System (CeNGEN) Project which is made possible by the support of NINDS R01NS100547.

## Author contributions

C.A. Davison, M. Hammarlund, and E.M. Jorgensen conceived the study. C.A. Davison performed and analyzed all experiments in this study. D. Garcia helped generate many of the strains used in this study and quantified axon breakage in multiple genotypes. C. Feng performed cloning, confocal microscopy, and provided technical advice. H. Hao generated the XE3230 strain used in this study and provided technical advice. The manuscript was written by C.A. Davison and M. Hammarlund, with input from all coauthors.

## Declarations of Interests

The authors declare no competing interests.

---

**Figure S1:**
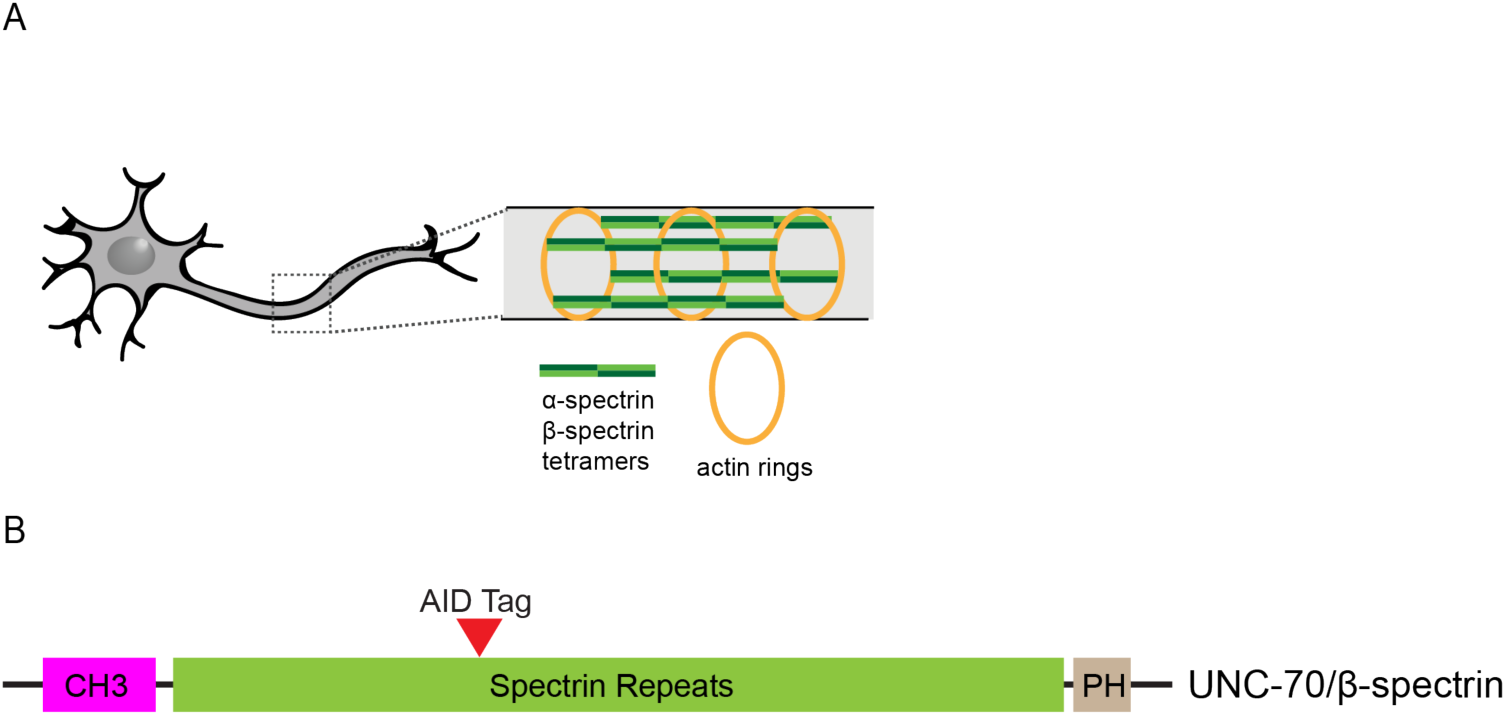
Schematic of the MPS and protein domain structure of *C. elegans* UNC-70/β-spectrin. (A) Cartoon of the MPS comprised of α-spectrin and β-spectrin tetramers linking actin rings (B) Schematic of the protein domain structure of UNC-70/β-spectrin and location of the AID tag

**Figure S2:**
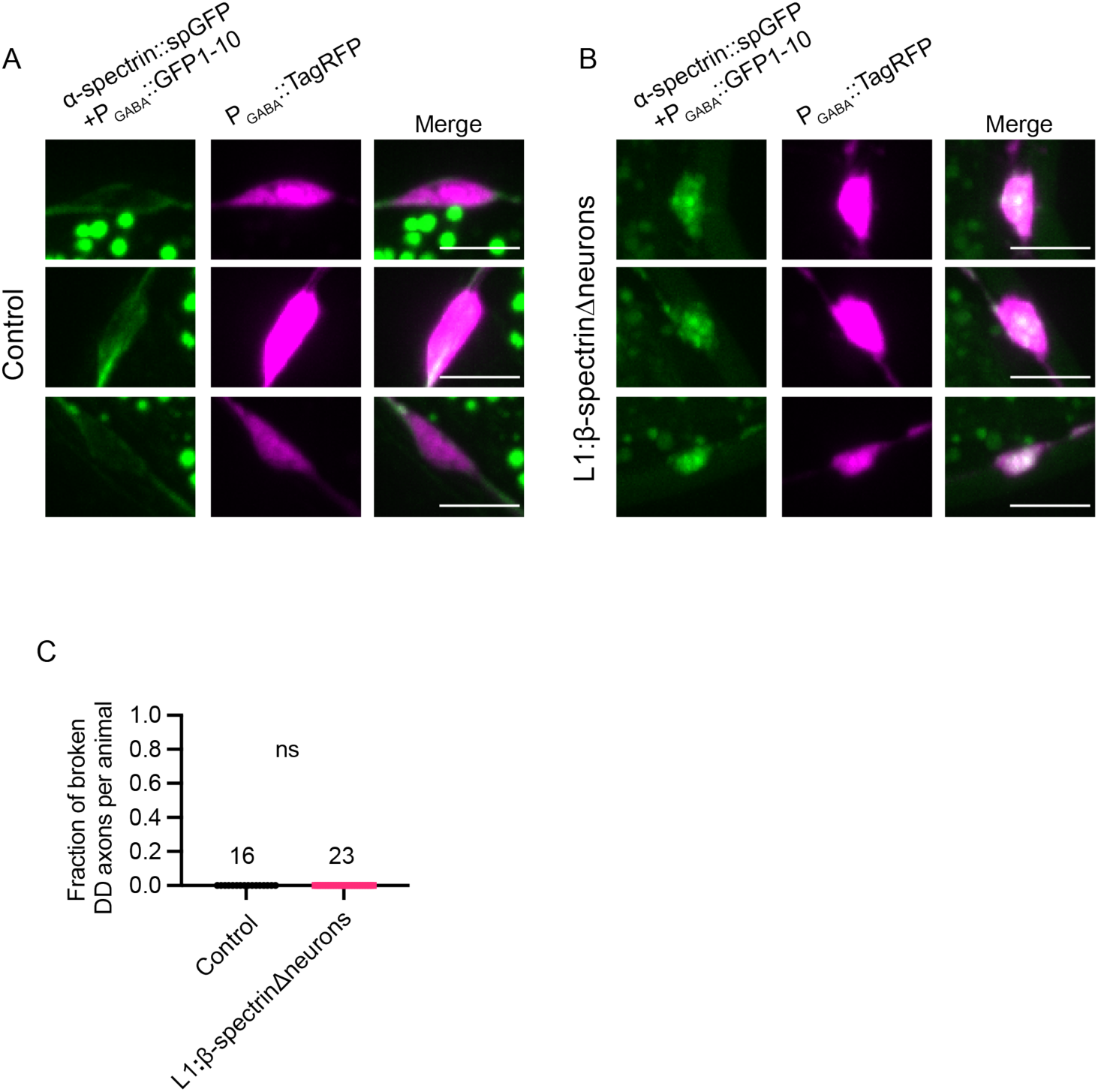
Degradation of neuronal β-spectrin for 24 hours results in accumulation of α-spectrin in DD cell bodies but unbroken DD commissures at the L1 stage. Representative images of endogenous α-spectrin (split-GFP) in DD cell bodies (TagRFP) in (A) control and (B) L1:β-spectrinΔneurons 24 hours after egg lay reveals accumulation of α-spectrin in cell bodies when the MPS is disrupted by degradation of neuronal UNC-70/β-spectrin. Scale bar 5μm. (C) Quantification of the fraction of broken DD axons 24 hours post egg-lay onto control plates or 1mM auxin (L1:β-spectrinΔneurons). The number of animals scored is shown above each dataset. All animals had 5 visible and intact DD commissures.

**Figure S3:**
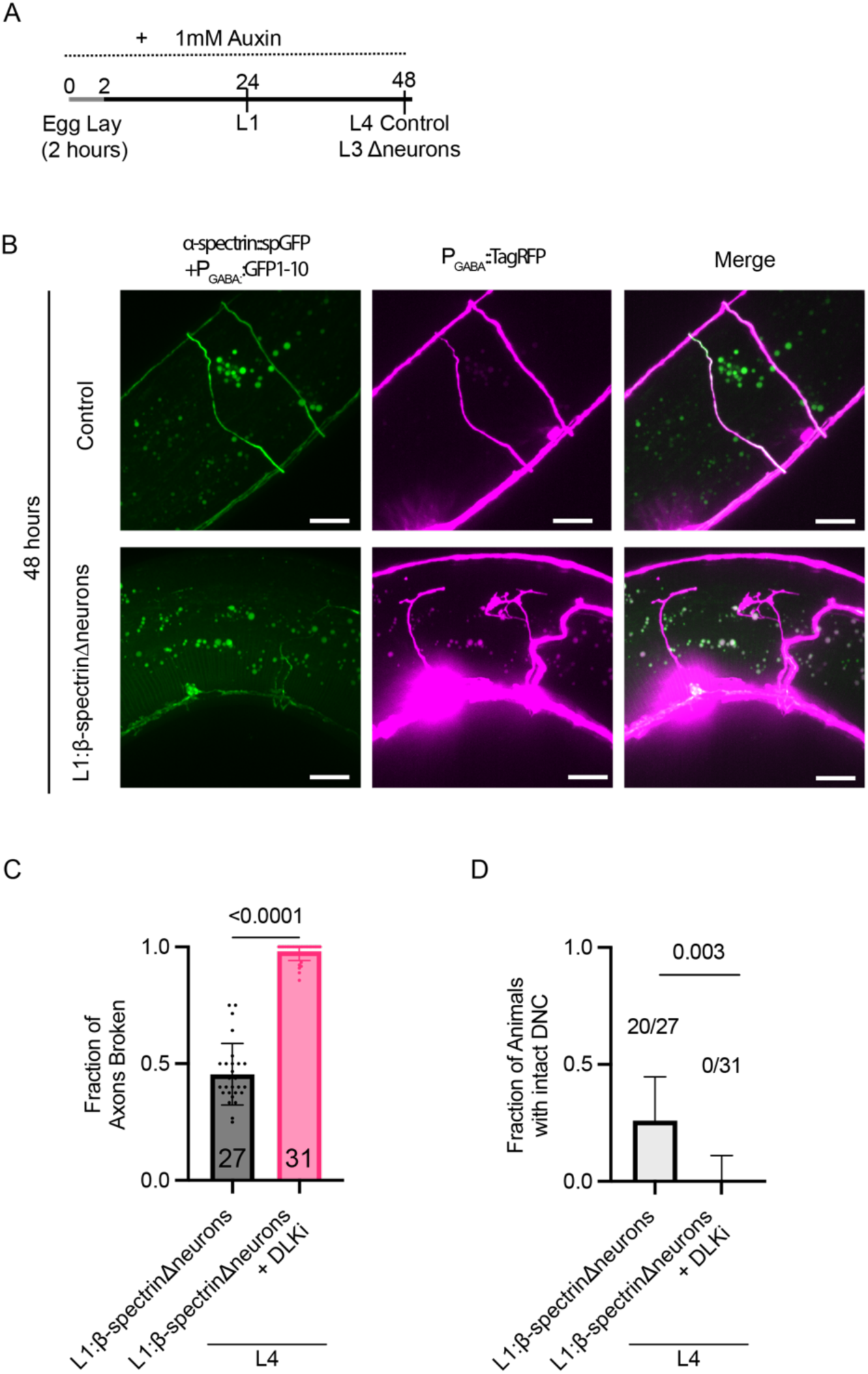
Axons break and regenerate when neurons lack β-spectrin. (A) Schematic of degradation timing for L1:β-spectrinΔneurons assayed at the 48 hour time point. At this timepoint, control animals are L4 stage while L1:β-spectrinΔneurons are slightly developmentally delayed (late L3/early L4) (B) Representative images of endogenous α-spectrin in GABA neurons (split-GFP labeled) and GABA neuron morphology (cytosolic TagRFP) in L4 control and L3/L4 L1:β-spectrinΔneurons animals exposed to 1mM Auxin for 48 hours. Scale bar 10 μm. Extensive growth cone formation is evident in L1:β-spectrinΔneurons animals. (C) Quantification of the fraction of axons broken per animal in L4 stage L1:β-spectrinΔneurons animals with and without inhibition of DLK-1 (DLKi, 50μM GNE-3511) (unpaired two-tailed t-test: p<0.0001). (D) Quantification of the fraction of animals with an intact dorsal nerve cord (DNC) in L4 stage L1:β-spectrinΔneurons animals with and without DLKi (Fisher’s exact test: p<0.0001).

**Figure S4:**
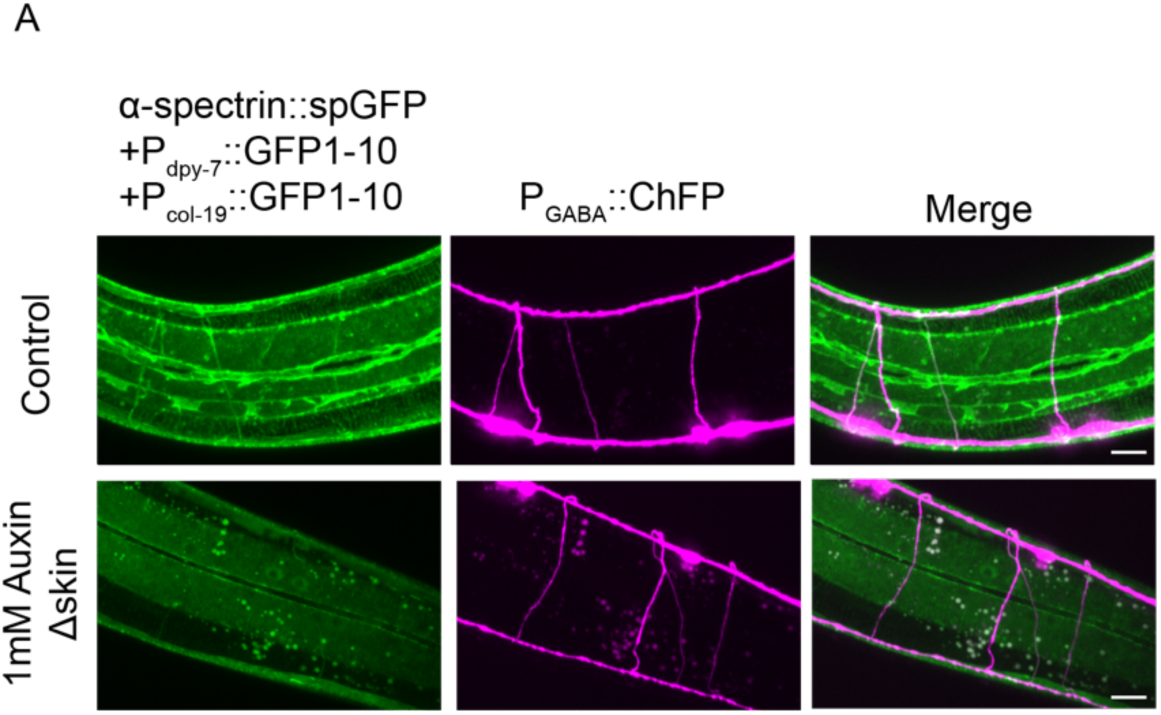
Degradation of epidermal β-spectrin successfully disrupts epidermal α-spectrin. Representative images of endogenous epidermal α-spectrin in control and L1:β-spectrinΔskin animals at L4 stage. Scale bar 10μm.

**Figure S5:**
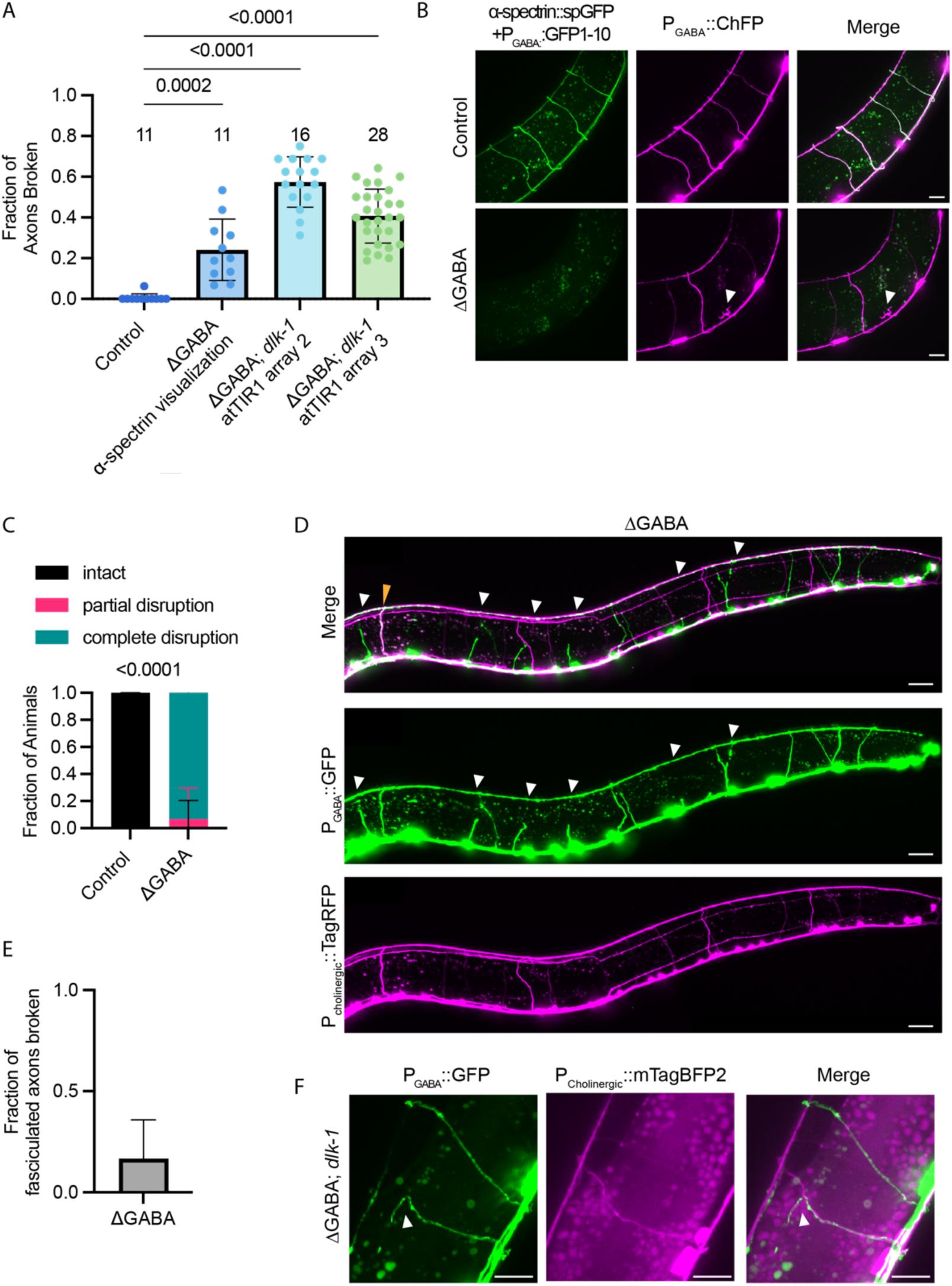
Neither variability of atTIR1 array nor direct scaffolding by cholinergic neuron commissures are responsible for the reduced breakage in L1:β-spectrinΔGABA animals. (A) Quantification of the fraction of axons broken in three additional array lines driving atTIR1 in GABA neurons under the *unc-47* promoter. The dataset labeled ΔGABA α-spectrin visualization is shown in panel B. Arrays 2 and 3 are in the *dlk-1* (j*u476*) background to prevent axon regeneration. (One-way ANOVA followed by Tukey’s multiple comparisons test p<0.0001). (B-C) Representative images and quantification of α-spectrin distribution in GABA neurons when β-spectrin is degraded from GABA neurons only. White arrow indicates broken axon. Scale bar 10μm. (Fisher’s exact test: p<0.0001). (D) Representative images of cholinergic (*P_unc-17_::TagRFP*) and GABAergic (*P_unc-47_::GFP*) neuronal commissures when β-spectrin is degraded from GABA neurons reveals scaffolding from neighboring cholinergic neurons is not common (only 1-2 events per animal). White arrows indicate broken VD/DD GABA axons. Yellow arrow indicates intact GABA commissure fasciculated with cholinergic neuron commissure. This fasciculation event occurs in most animals, and this axon has not been observed to break. Scale bar 20 μm (E) Quantification of the fraction of fasciculation events in which the VD/DD commissure is broken (4/24 broken). (F) Representative images of cholinergic (*P_unc-17_::mTagBFP2*) and GABAergic (*P_unc-47_::GFP*) neuronal commissures when β-spectrin is degraded from GABA neurons in the *dlk-1* background reveals fasciculation is not sufficient to prevent breakage. White arrow indicates broken VD/DD commissure. Scale bar 10 μm.

**Figure S6:**
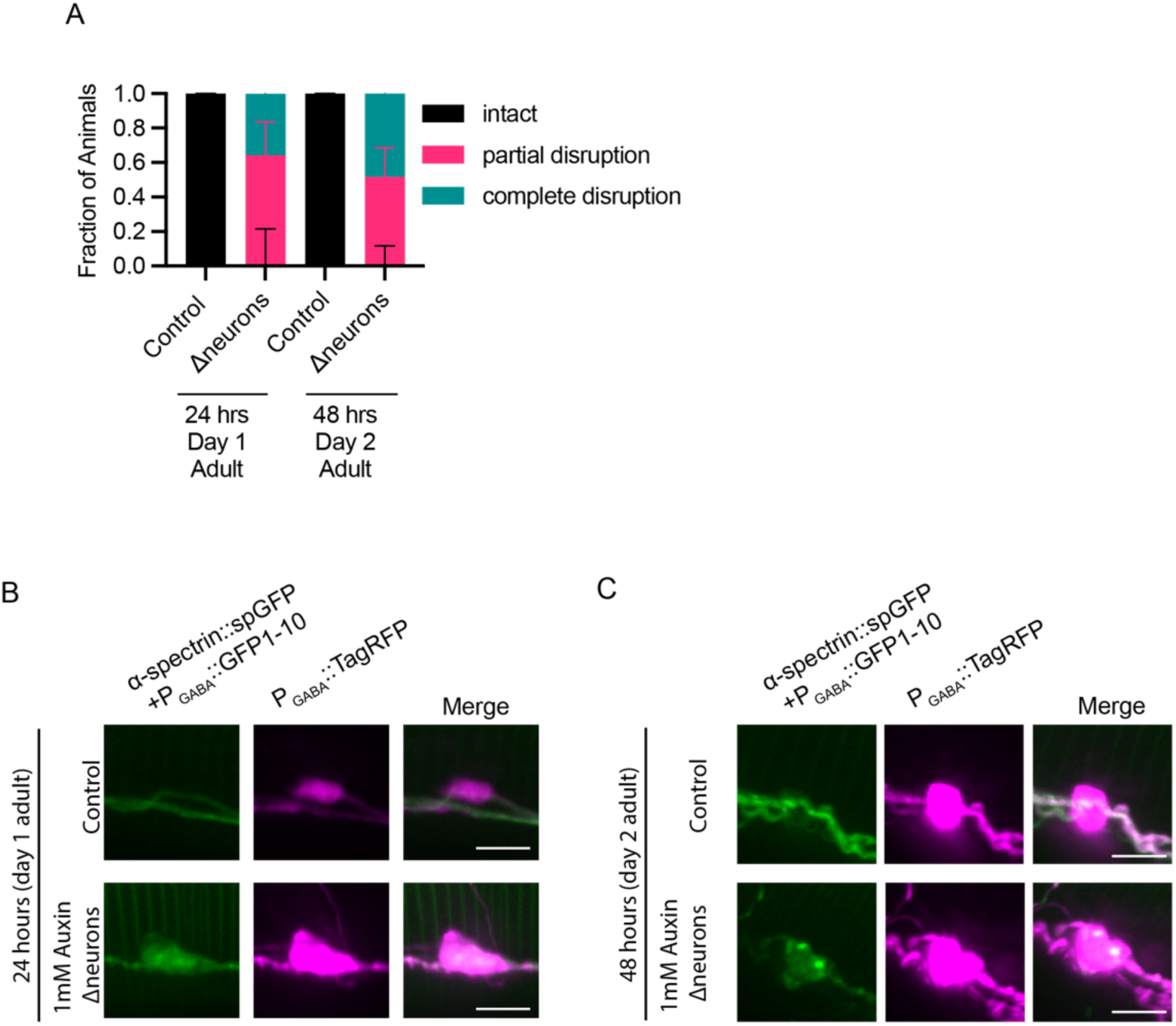
Degradation of β-spectrin in mature neurons disrupts the MPS. (A) Quantification of the fraction of animals with altered α-spectrin distribution in GABA neurons when neuronal β-spectrin is degraded for 24 or 48 hours starting at the L4 stage. Complete disruption of α-spectrin is defined as full loss of endogenous *spc-1* split-GFP signal in GABA neuron commissures. Partial disruption is defined as some split-GFP signal remaining in one or more commissures in the worm, but the signal is non-uniform. (B-C) Representative images of accumulation of α-spectrin (split GFP) in VD/DD cell bodies (magenta cytosolic TagRFP) upon neuronal β-spectrin degradation for 24 or 48 hours starting at the L4 stage. Scale bar 5 μm.

**Table S1:**
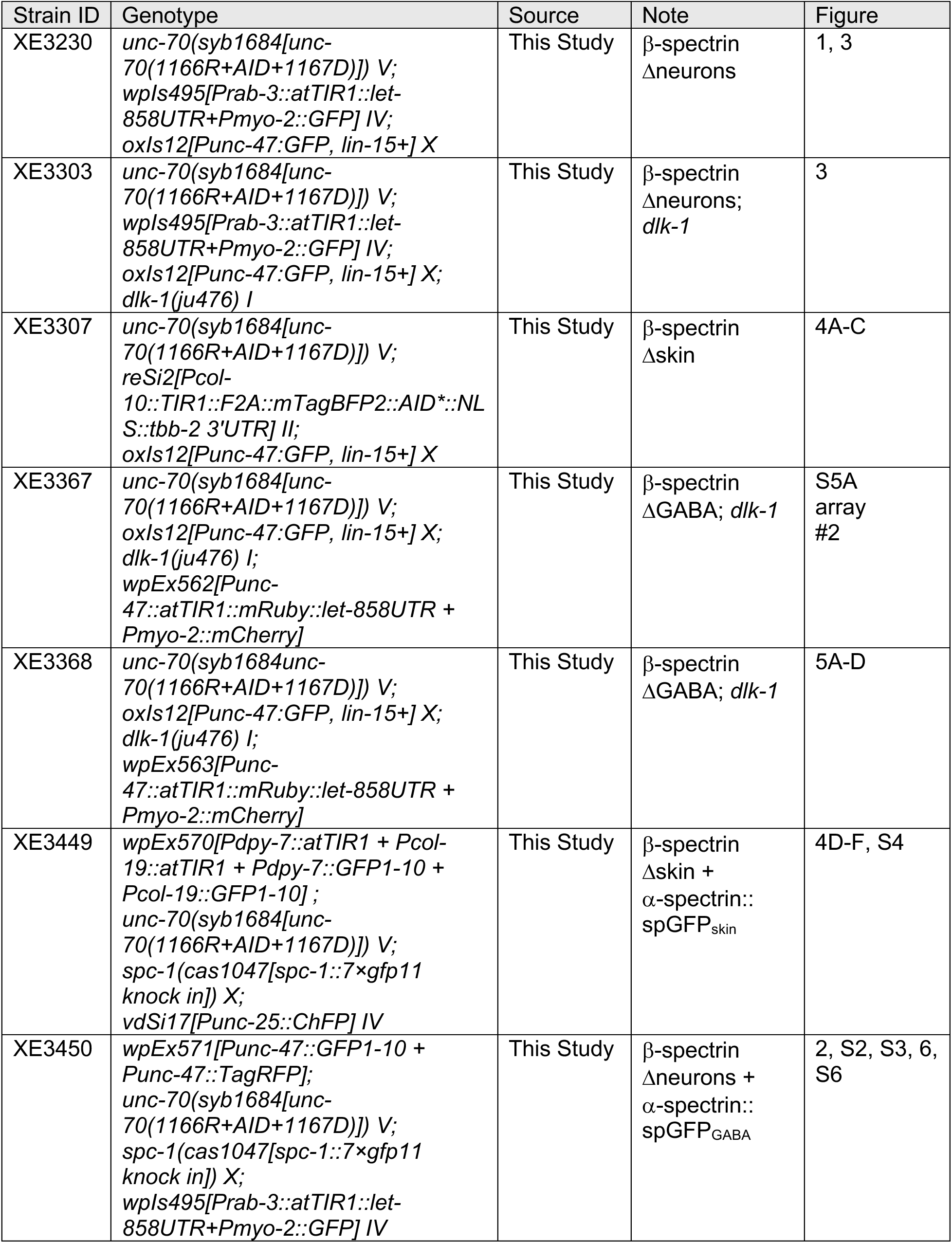
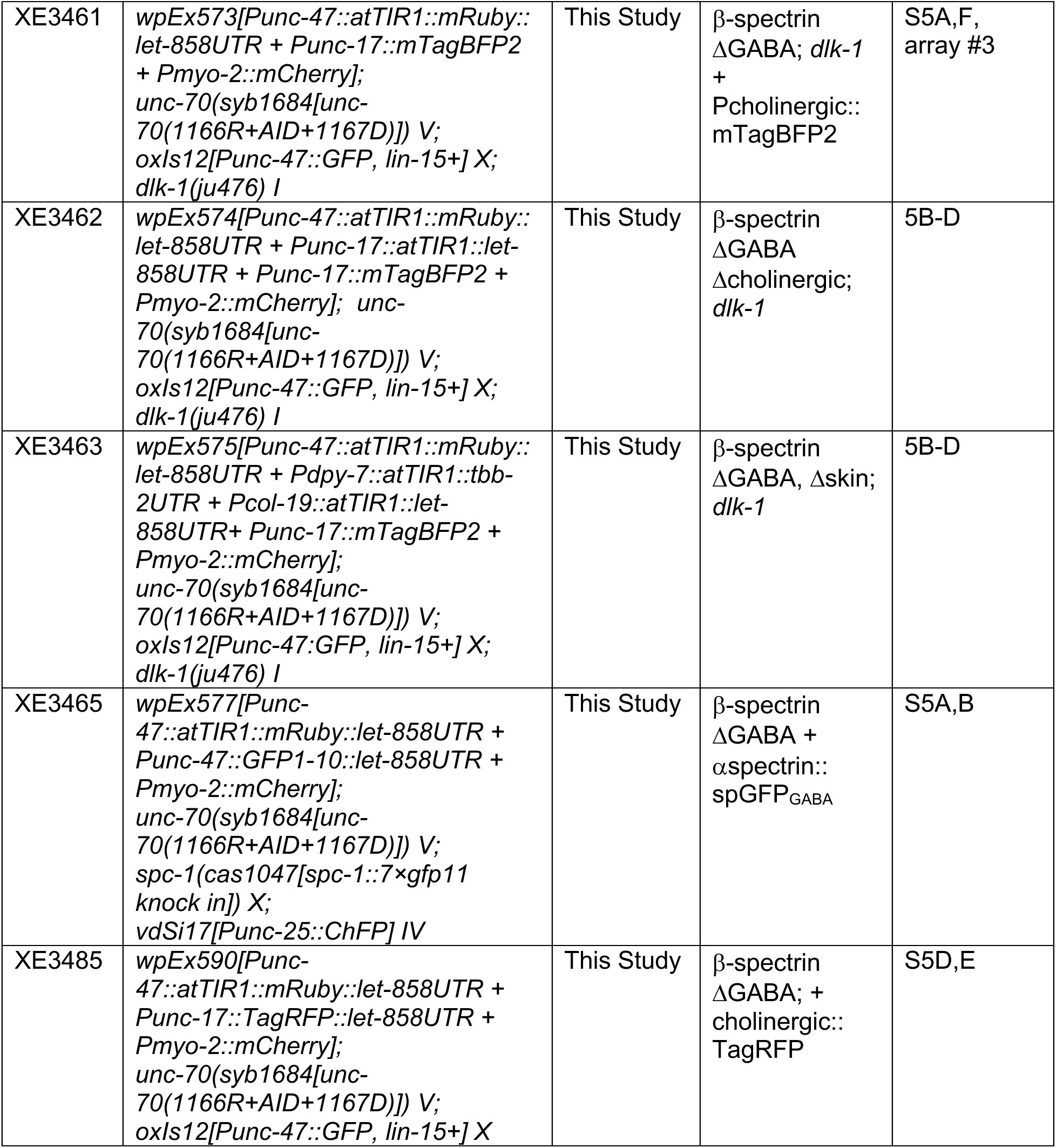
List of *C. elegans* strains used in this study.

**Table S2:** List of plasmids used in this study.

| Plasmid identifier | Description | Concentration Injected (ng/uL) |
| --- | --- | --- |
| pHHW-004 | Prab-3::atTIR1::let-858UTR | 20 |
| pPD118.33 | Pmyo-2::GFP | 5 |
| pSEM319 | Fragment of unc-119 MoSTI site Addgene: #159823 | 5 |
| pCFJ90 | Pmyo-2::mCherry | 1 |
| pCAD145 | Punc-47::atTIR1::mRuby::let-858UTR | 20 |
| pCAD165 | Pdpy-7::atTIR1::tbb-2UTR | 20 |
| pCAD154 | Pcol-19::atTIR1::let-858UTR | 20 |
| pCAD164 | Pdpy-7::GFP1-10::tbb-2UTR | 20 |
| pCAD163 | Pcol-19::GFP1-10::tbb-2UTR | 20 |
| pCAD136 | Punc-47::GFP1-10::let-858UTR | 20 |
| pCAD169 | Punc-47::TagRFP::let-858UTR | 20 |
| pCAD174 | Punc-17::mTagBFP2::let-858UTR | 20 |
| pCAD170 | Punc-17::atTIR1::let-858UTR | 20 |
| pCAD179 | Punc-17::TagRFP::let-858UTR | 30 |

## Notes

### Competing Interest Statement

The authors have declared no competing interest.

